# Horizontal gene transfer of the *pirAB* genes responsible for Acute Hepatopancreatic Necrosis Disease (AHPND) turns a non-*Vibrio* strain into an AHPND-positive pathogen

**DOI:** 10.1101/2019.12.20.884320

**Authors:** Sarmila Muthukrishnan, Tom Defoirdt, Mohamed Shariff, Ina-Salwany M. Y, Fatimah Md Yusoff, I. Natrah

**Affiliations:** Department of Aquaculture, Faculty of Agriculture, Universiti Putra Malaysia, 43400 UPM, Serdang Selangor; Laboratory of Marine Biotechnology, Institute Bioscience, Universiti Putra Malaysia, 43400 UPM, Selangor; International Institute of Aquaculture and Aquatic Sciences, Universiti Putra Malaysia, 43400 UPM, Serdang, Selangor; Center for Microbial Ecology and Technology, Ghent University, Coupure Links 653, 9000 Gent, Belgium

## Abstract

In the past decade, shrimp farms, particularly those established in Asia, Mexico and South America suffered from the outbreak of an emergent penaeid shrimp disease known as Acute Hepatopancreatic Necrosis Disease (AHNPD). The PirA and PirB toxins produced by plasmid pVA1 in *Vibrio parahaemolyticus* were reported to cause the AHPND pathology. More recent research demonstrated that *V. parahaemolyticus* is not the only species that can cause AHPND, as other *Vibrio* species were also found to contain PirAB-containing plasmid. The present study assessed the Horizontal Gene Transfer (HGT) of AHPND that transforms genes (*pirA* and *pirB*) from AHPND positive *V. parahaemolyticus* to non-AHPND and non-vibrio species identified as *Algoriphagus* sp. strain NBP. The HGT of *pirA* and *pirB* genes from the AHPND positive *V. parahaemolyticus* to *Algoriphagus* sp. strain NBP was found to occur at different temperatures. The conjugation efficiency rate (n°) of *pirAB* from *V. parahaemolyticus* to *Algoriphagus* sp. strain NBP at 30°C and 40°C showed 80-91% efficiency. Shrimp challenged with the *pirA* and *pirB* positive *Algoriphagus* sp. strain NBP also demonstrated typical pathognomonic AHPND lesions during the histopathologic examination.

**Author summary:** AHPND is a significant threat to the shrimp industry leading to high losses. The results demonstrated that the conjugative transfer of the *pirA* and *pirB* positive *V. parahaemolyticus* (donor strain) to a non-*Vibrio* and non-pathogenic bacterium (recipient strain), successfully transformed the non-pathogenic bacterium into a disease-causing strain with a disease-causing capability similar to the donor strain. Initially, *V. parahaemolyticus* that express the PirA and PirB toxins which encoded by a conjugative plasmid cause sloughing and degeneration of shrimp hepatopancreatic.

## Introduction

It has been a decade since the emergence of Acute Hepatopancreatic Necrosis Disease (AHPND) that has caused global losses of more than $ 1 billion per year, and mortality rates up to 100% in the shrimp farming industries within the regions of Asia, Mexico, South America and Texas [1–2]. The etiological agent has been initially identified as *Vibrio parahaemolyticus* carrying a plasmid containing toxin genes (*pirA* and *pirB*). The two toxin subunits, PirA and PirB, are homologous to the Pir (*Photorhabdus* insect-related) binary toxin. Recent studies have reported that other vibrios that are closely related to *Vibrio parahaemolyticus*, such as *Vibrio campbellii* from Vietnam [3], *Vibrio owensii* from China [4], *Vibrio campbellii* from China [5], *Vibrio punensis* from South America [6] and *Vibrio harveyi* from Malaysia [7] also demonstrated AHPND pathology in shrimp. The presence of conjugative transfer genes on the pVA1 plasmid (70 kb plasmid harboring the *pirA* and *pirB* genes) [8] postulates the possibility of mobilisation of AHPND virulence genes to other *Vibrio* spp. via horizontal gene transfer [9].

The presence of both *pirA* and *pirB* genes in other *Vibrio* spp. suggested that the toxin genes are transmissible through conjugal transfer, thereby turning the acceptor bacterium into an AHPND-causing strain [8]. This has significant implications for disease management, as the presence of non-pathogenic *Vibrio* strains that are closely related to *Vibrio parahaemolyticus* also includes risk as these bacteria might be transformed into pathogens through horizontal gene transfer. Indeed, *pirA* and *pirB* genes have been well characterised in many *Vibrio* spp. with AHPND-like histopathology [3, 5, 7]. Thus far, all *pirA* and *pirB* positive strains which causes AHPND belong to species within the Harveyi clade of vibrios (i.e. a clade of species closely related to *V. harveyi*) [3–7]. However, very little is known on the possibility of conjugative transfer of these genes to non-*Vibrio* spp. Such a gene transfer would have far-reaching implications for disease control as it would be able to also turn non-*Vibrio* strains into AHPND-causing agents. In order to investigate potential horizontal gene transfer of the *pirA* and *pirB* genes from *V. parahaemolyticus* to a non-AHPND and non-*Vibrio* bacterium, we assessed transfer of the genes to an *Algoriphagus* sp. strain NBP isolated from marine microalgae (*Nannochloropsis* sp.).

## Results

### Isolation and of a pirAB negative non-*Vibrio* strain from a microalgal culture

Eight strains were isolated from the microalgal culture. An isolate with a pink colony and creamy texture on MA plates, denoted NBP, was selected for further work as this colony morphology enabled us to easily differentiate the isolate from *Vibrio parahaemolyticus* BpShHep31 (Fig 1). The selected isolate was confirmed to be negative for the *pirA* and *pirB* genes using PCR with specific primers (VpPirA-284F, VpPirA-284R, VpPirB-392F, and VpPirB-392R).

**Fig 1.**
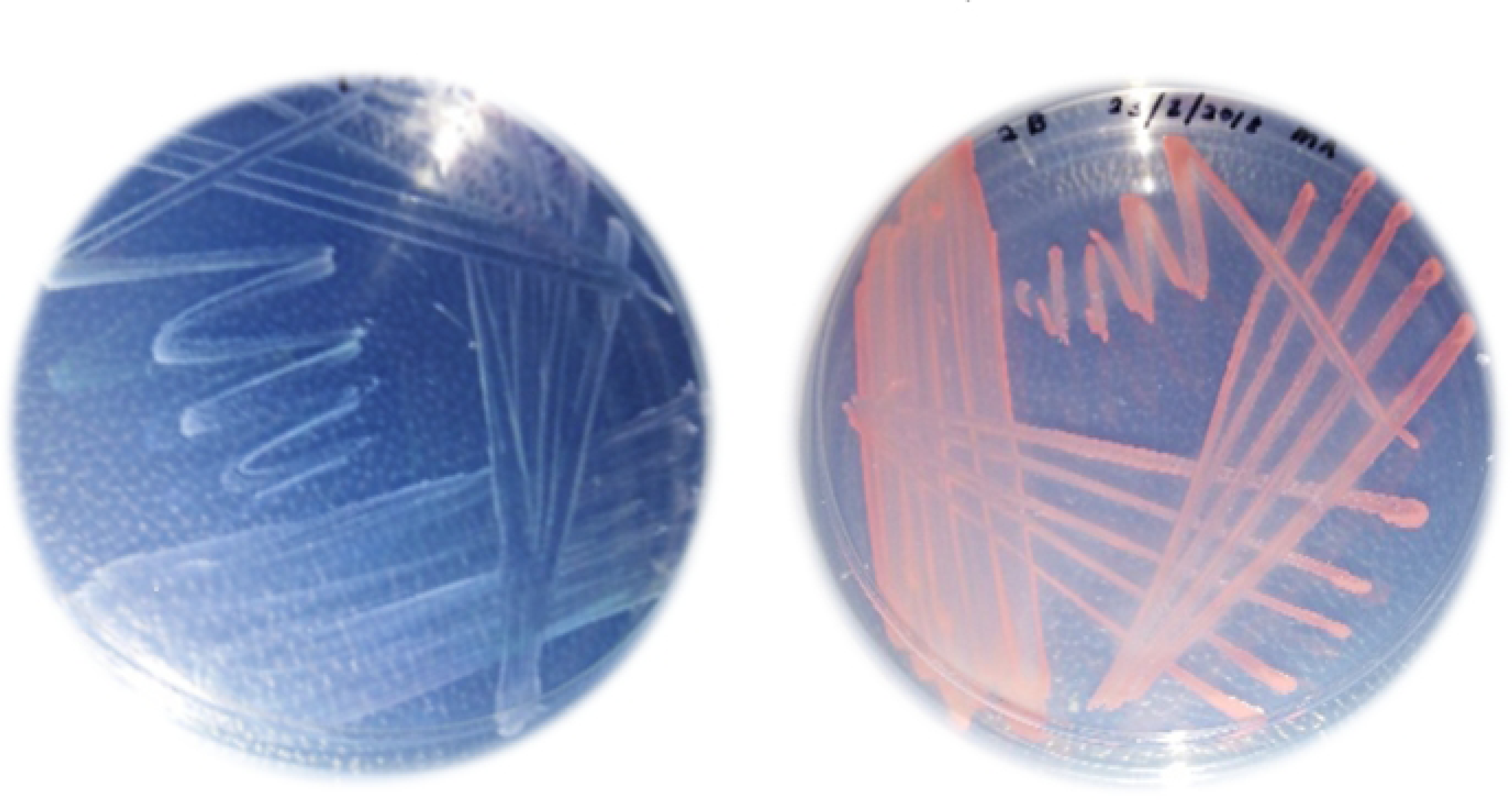
Pictures of *Vibrio parahaemolyticus* BpShHep31 (left) and strain NBP isolated from a microalgal culture (right) on Marine Agar plates showing the clearly distinct colony morphologies.

### Identification of the pirAB negative isolate

A BLAST search revealed that the 16S rDNA gene sequence of isolate NBP showed 99% similarity to that of *A. marincola* strain SW-2 (GenBank accession **MK583623**). The 16S rDNA of the isolate formed a monophyletic taxon with *A. marincola* strain SW-2 with a posterior probability (PP = 0.67) (Fig 2). Hence, the isolate is further denoted as *Algoriphagus* sp. strain NBP. The 16S rDNA sequence from strain NBP has been submitted to GenBank under accession number **MK583623**.

**Fig 2.**
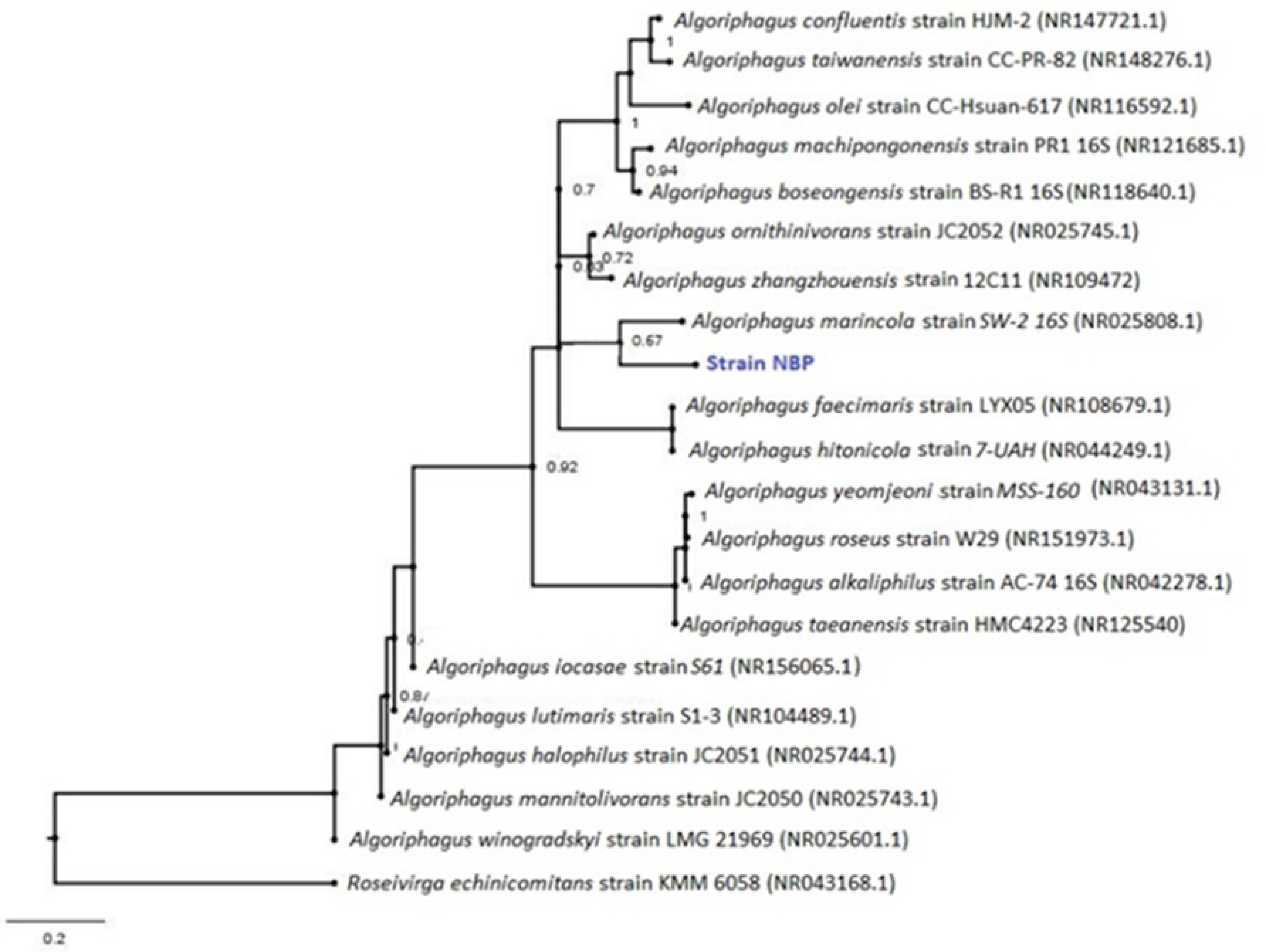
Phylogenetic reconstruction of *Algoriphagus* sp. strain NBP based on 16S rDNA sequences using mixed model method. Percentage bootstrap values (10,000,000 replicates) > 65% are presented.

### Co-culture of isolate NBP and *V. parahaemolyticus* BpShHep31 and screening for the presence of pirAB genes in colonies re-isolated after co-culture

The results showed that although *Algoriphagus* sp. strain NBP was negative for *pirA* and *pirB* prior to co-culture, several colonies of the isolate that were picked up from MA plates after co-culture with *Vibrio parahaemolyticus* BpShHep31 tested positive for the presence of *pirA* and *pirB* (Fig 3). Both *PirA* (GenBank accession no. **MN652913**) and *PirB* (GenBank accession no. **MN652914**) sequences from *Algoriphagus* sp. strain NBP demonstrated 99% similarities when compared to *pirAB* genes from *V. parahaemolyticus* BpShHep31.

**Fig 3.**
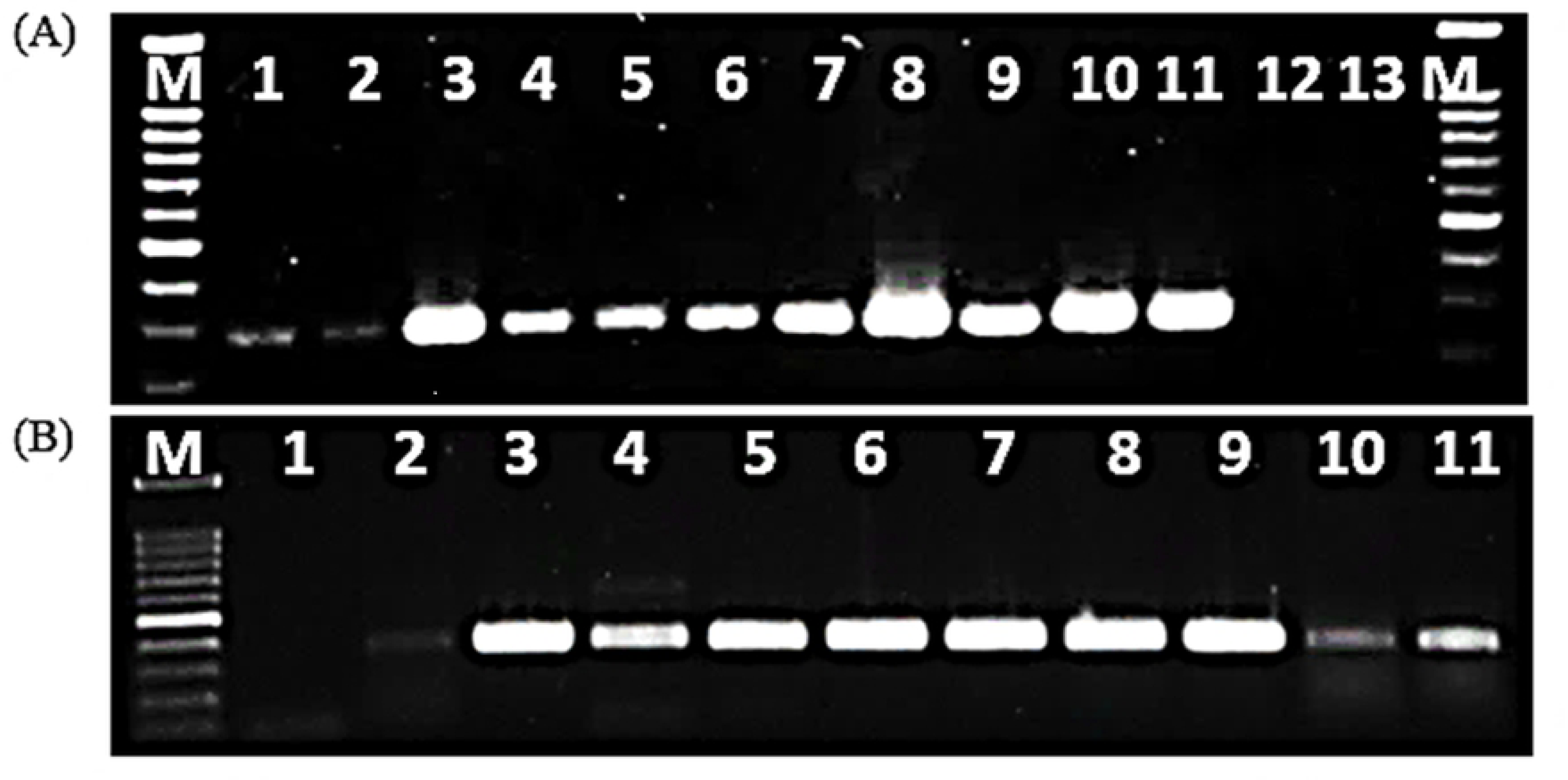
Gel electrophoresis of PCR products. (**A**) Gel electrophoresis of PCR products after amplification of the *pirA* gene from *Algoriphagus* sp. strain NBP colonies picked up after co-culture with *V. parahaemolyticus* BpShHep31. Lanes 1-3: triplicates at 20 °C, lanes 4-6: triplicates at 30 °C, lanes: 7-9: triplicates at 40 °C, lanes 10-11: positive control (*V. parahaemolyticus*, BpShHep31), lanes 12-13: negative control, M: 100 kb ladder. (**B**) Gel electrophoresis of PCR products after amplification of the *pirB* gene from *Algoriphagus* sp. strain NBP colonies picked up after co-culture with *V. parahaemolyticus* BpShHep31. Lane 1: negative control, lane 2: positive control, lanes 3-5: triplicates at 20 °C, lanes 6-8: triplicates at 30 °C and lanes 9-11: triplicates at 40 °C.

The density of *Algoriphagus* sp. strain NBP in the cocultures significantly increased (*P* < 0.05) at higher temperature (Fig 4). The *Algoriphagus* sp. strain NBP colonies isolated from the cocultures were subjected to screening for the presence of *V. parahaemolyticus toxR* in order to exclude contamination with *V. parahaemolyticus*. The results demonstrated that 90-92 % of the *Algoriphagus* sp. strain NBP colonies were negative for *toxR*, indicating that the contamination of *Algoriphagus* sp. strain NBP colonies with *V. parahaemolyticus* was less than 10%. Furthermore, the *Algoriphagus* sp. strain NBP incubated in the MB overnight and plated on TCBS showed no growth indicating the absence of *V. parahaemolyticus*.

**Fig 4.**
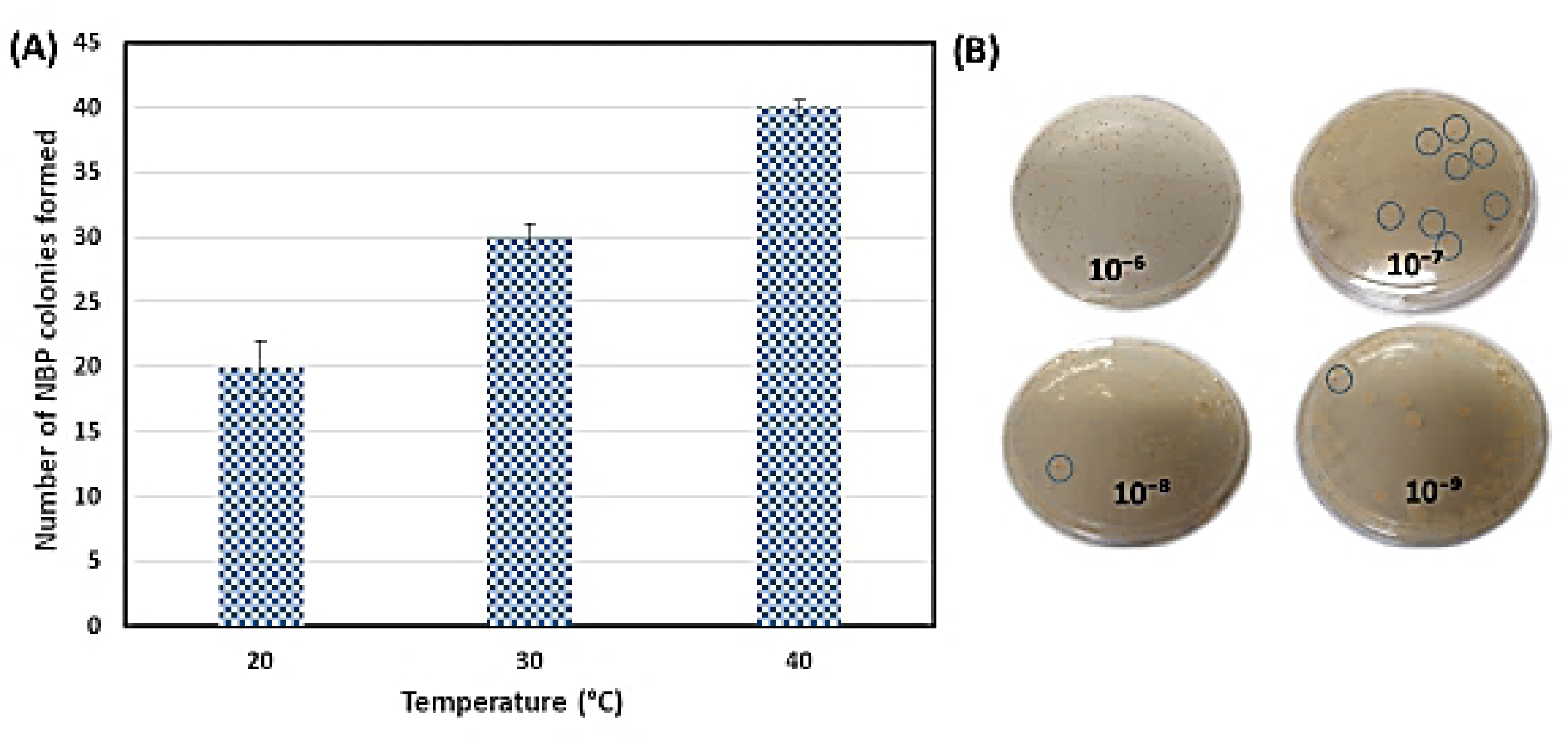
Number of *Algoriphagus* sp. strain NBP colonies. (A) Number of *Algoriphagus* sp. strain NBP colonies formed at 10^8^ CFU mL^−1^ upon co-culture. (B) Plate picture of the co-culture at different dilutions (from 10^−6^ to 10^−9^).

We calculated the conjugation efficiency of *pirAB* genes and found that it was high (80-81%) at 30°C and 40°C (Table 1). Furthermore, no significant differences (*P* > 0.05) were observed between the conjugation efficiency of *pirAB* at 30-40°C.

**Table 1.** Conjugation efficiency of the *pirA* and *pirB* genes from *V. parahaemolyticus* to *Algoriphagus* sp. strain NBP.

| Temperature (°C) | Conjugation efficiency (%) |  |
| --- | --- | --- |
|  | *pirA* | *pirB* |
| 20 | 68 ± 4<sup>a</sup> | 58 ± 7<sup>a</sup> |
| 30 | 81 ± 10<sup>b</sup> | 72 ± 4<sup>b</sup> |
| 40 | 80 ± 8<sup>b</sup> | 80 ± 9<sup>b</sup> |

<sup>ab</sup> Mean value (mean ± SD) with different superscript letters are significantly different ( $P < 0.05$ )

### Shrimp immersion challenge with the pirAB positive *Algoriphagus* sp. strain NBP

The pathogenicity of the *pirA* and *pirB* positive *Algoriphagus* sp. strain NBP was investigated through an *in vivo* immersion assay using *P. vannamei*. Shrimp cultures that were inoculated with the *pirA* and *pirB* positive *Algoriphagus* sp. strain NBP showed significant (*P* < 0.05) mortality when compared to unchallenged control cultures, whereas shrimp cultures that were challenged with the *pirA* and *pirB* negative *Algoriphagus* sp. strain NBP showed no significant mortality (Fig 5). Additionally, the shrimp challenged with *pirA* and *pirB* positive *Algoriphagus* sp. strain NBP demonstrated pale hepatopancreas, lethargy and lack of appetite.

**Fig 5.**
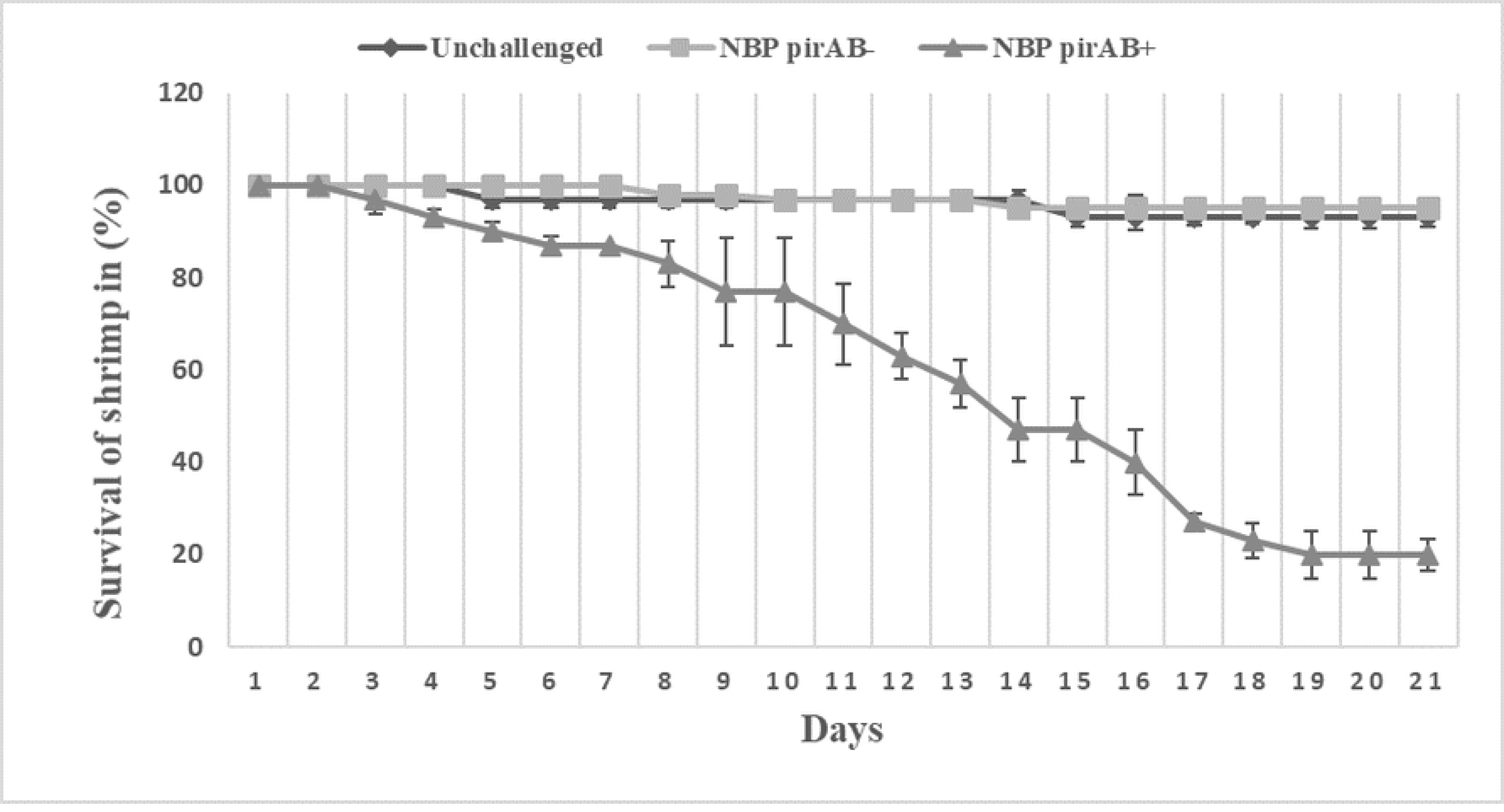
Survival of *P. vannamei*. Survival of *P. vannamei* without inoculation of any bacteria (Unchallenged), challenged with non-*PirAB* Algorhipagus sp. strain NBP (NBP *pirAB+*), and *pirAB* positive *Algoriphagus* sp. strain NBP (NBP *pirAB-*). The bacteria were inoculated to rearing water at the start of the experiment at 10^6^ CFU/mL. Error bars represent the standard deviation of triplicate shrimp cultures.

The shrimp challenged with *pirA* and *pirB* positive *Algoriphagus* sp. strain NBP demonstrated typical AHPND pathology on the 14^th^ day. Histopathologic evaluation revealed detachment of epithelial cells from the membrane up to lumen (see Fig 6A), which affected the integrity of the tubules. Loose hepatopancrease, tubule atrophy, as well as lack of B, R and F cells, were observed (see Fig 6B). The formation of hemocytic encapsulation and the massive sloughed hepatopancrease can be noted in Figs 6C and 6D. Overall, the typical pathognomonic AHPND lesions were observed in shrimp challenged with the *pirA* and *pirB* positive *Algoriphagus* sp. strain NBP. Expected *pirA* and *pirB* amplicons were also generated from DNA extracts of shrimp challenged with NBP *pirAB*+, whereas samples taken from unchallenged shrimp and shrimp challenged with NBP *pirAB-* tested negative.

**Fig 6.**
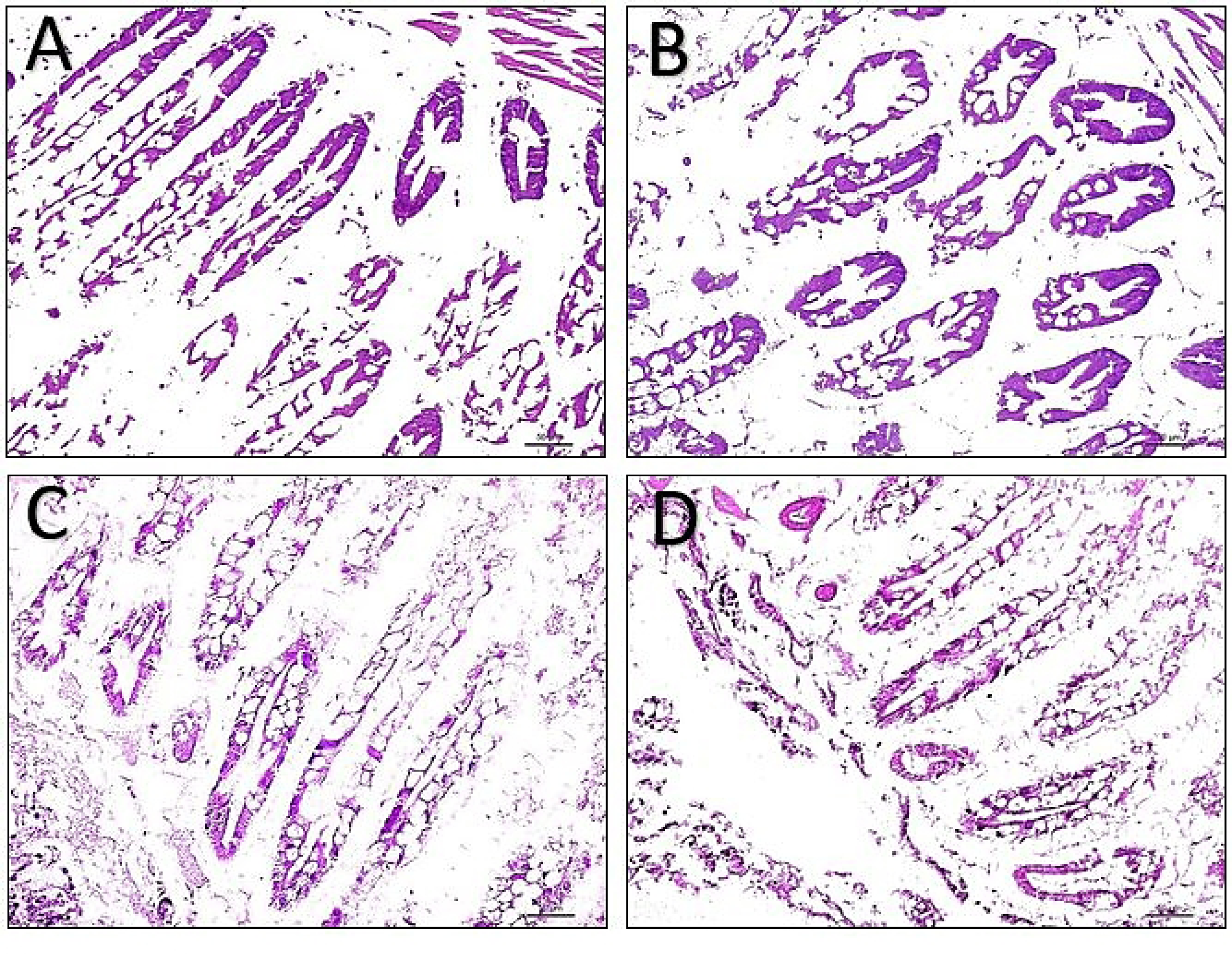
Photomicrographs of shrimp hepatopancreas. Photomicrographs of shrimp hepatopancreas challenged with *pirA* and *pirB* positive *Algoriphagus* sp. strain NBP at 10^6^ CFU mL^−1^ (at 14^th^ day of challenge). (A) Histological sections show tubule atrophy and detachment of epithelial cells. (B) Sloughing of cells to lumen and degeneration of cells were observed. (C) Hemocytic infiltration and massive sloughing of hepatopancreas recorded. (D) Histological sections show severe necrosis, melanized hemocytic nodules, tubule atrophy, elongated lumen and karyomegaly. Scale bars: 50 µm, magnification: 20 x.

## Discussion

The *V. parahaemolyticus* strain BpShHep31 used in this study were previously isolated from diseased *P. vannamei* [7] and were tested positive for *pirAB* genes using VpPirA and VpPirB primer sets [10]. Many studies conducted HGT experiment using mutant strains to determine the conjugation activity and efficiency [11–13]. In this study, we tested on phenotypically different bacterium isolated from marine microalga *Nannochloropsis* sp. which was previously isolated from a shrimp pond.

The possibility of AHPND toxin genes to spread via conjugation and permanently be inheritable in the recipient bacteria has been highlighted by Lee et al. [8]. The previous study demonstrated that *V. parahaemolyticus* strain M2-36 contain gene fragments that are flanked by transposase coding sequence, which is also known as mobile genetic elements (MGEs). The presence of MGEs in *V. parahaemolyticus* strain M2-36 suggested the acquisition or deletion of *PirA* and *PirB* might be due to HGT [8]. Hence, to prove that HGT could also occur between a *Vibrio* and a *non-Vibrio* strain, a co-culture experiment between AHPND positive *Vibrio parahaemolyticus* BpShHep31 with a non-AHPND and non-vibrio bacterium identified as *Algorhipagus* sp. strain NBP was conducted. In this study, three different temperatures were evaluated to observe the impact of temperature on HGT among the different bacteria. The results demonstrated that HGT occurred in all the three incubation temperatures (20 °C, 30 °C, and 40 °C) and time points (24 h, 48 h, and 72 h). The *V. parahaemolyticus* strain BpShHep31 functioned as a donor by donating both *pirA* and *pirB* containing plasmid to genes to the recipient species (*Algorhipagus* sp. strain NBP) during the co-culture experiment. To the best of the authors’ knowledge, this is the first study on the induction of *pirA* and *pirB* genes via HGT from AHPND *Vibrio* to a non-*Vibrio* strain.

Thus, to further investigate the expressions of *pirA* and *pirB* genes in the recipient cells, the successful recipient cells were sub-cultured in an *in vivo* challenge test. Interestingly, shrimp challenged with the recipient cells (*Algorhipagus* sp. strain NBP) carrying *pirA* and *pirB* virulence genes exhibited AHPND pathology and demonstrated significant mortalities (*P* < 0.05). However, *V. parahaemolyticus* strain BpShHep31 showed rapid mortalities in *P. vannamei* shrimp (50% mortalities within 2 days) [7] compared to the recipient cells (*Algorhipagus* sp. strain NBP) with the *pirAB* genes. Recent study also reported on the presence of *pirA*- and *pirB*-like genes in a non-vibrio *Micrococcus luteus* [14]. However, there is no data on the *in vivo* challenge test and the mechanism by which *Micrococcus luteus* obtained the *pirAB* genes.

Despite the genomic content in the plasmids of AHPND strains isolated from Asia, Mexico, and South America seemed to be distinct [10], all AHPND strains have been proven to carry a group of transposase-coding sequence linked to HGT [15]. Han et al [15] also discovered nine proteins in AHPND positive *Vibrio* spp., which were identical to proteins encoded with ORFs involved in plasmid conjugation and mobilisation. Besides, Dong et al., 2019 demonstrated the conjugation transfer of *pirAB* genes from *V. parahaemolyticus* strain *Vp*2S01 to non-AHPND *V. campbellii* with a transfer efficiency of 2.6×10^−8^ transconjugant/recipient. This study demonstrated 80% of transfer efficiency from *V. parahaemolyticus* strain BpShHep31 to non-*Vibrio* recipient bacteria (*Algorhipagus* sp. strain NBP) at 30-40°C. This finding further justifies that both *pirA* and *pirB* genes could be transferred to the non-pathogenic and non-*Vibrio* recipient bacteria (*Algorhipagus* sp. strain NBP) through conjugation. The transfer of *pirA* and *pirB* genes from pathogenic bacteria to non-pathogenic bacteria may then contribute to the emergence of new AHPND strains.

In summary, the study data demonstrated that *pirA* and *pirB* genes can be easily transferred to other microbes (non-*Vibrio* bacteria). The study outcomes postulate the presence of multiple non-*Vibrio* species in the shrimp ponds with *pirA* and *pirB* genes. This study shows that future researches should also consider non-*Vibrio* bacteria for AHPND screening process. The rapid intraspecies and interspecies HGT of *pirA* and *pirB* genes increased the complication for identifying the causative agents of AHPND. This scenario catalyses our need to determine preventive and mitigation measures to curb the spread of these pathogenic genes. Furthermore, comprehending the mechanism of interspecies gene transfer between AHPND *Vibrios* and non-AHPND, non-*Vibrio* bacterium is indeed crucial in light of this shrimp pandemic.

## Methods

### Bacterial strains and culture conditions

Acute hepatopancreatic necrosis disease (AHPND)-positive *Vibrio parahaemolyticus* strain BpShHep31 [7] was used as a donor of the *pirAB* containing plasmid. Ten microliters of a stored culture of the strain in 40% glycerol at −80 °C were plated onto Marine Agar (MA) (Difco™ overnight at 28 °C. A single colony was picked from the plate and cultured in MB at 28 °C under constant agitation (150 rpm).

### Isolation of bacteria from *Nannochloropsis* sp

The marine green microalga *Nannochloropsis* sp. used in this study was obtained from Bioproduct Lab (BP), University of Putra Malaysia (UPM), Malaysia. The microalga was cultured in Guillard’s f/2 medium at room temperature in 100 µmol of photons m^−2^s^−1^ of light intensity and 30 ppm of salinity. One mL of the culture was transferred to a 1.5 ml Eppendorf tube and centrifuged at 3000 rpm for 5 minutes. Next, 100 µL of the supernatant was serially diluted using saline buffer and plated on MA. The plates were incubated overnight at 28 °C. Different colonies were picked based on their morphology, colour, and structure on the MA plates. The isolated colonies were sub-cultured twice on MA and were subsequently cryopreserved at −80 °C in Marine Broth (MB) (Difco™, USA) containing 40% of glycerol. A single isolate, denoted NBP, was selected for further experiments based on its colony morphology being clearly different from that of *Vibrio parahaemolyticus* BpShHep31 which thus enabled easy differentiation after co-culture experiments.

### Screening for the presence of *pirA* and *pirB* genes

The presence of *pirA* and *pirB* genes was determined using PCR with specific primers. Genomic DNA was extracted from grown cultures using the Geneaid kit (Taiwan) and was purified by adhering to the protocol provided by the manufacturer. The primers used for PCR amplification of the *pir* genes are listed in Table 2. The PCR mixtures were composed of 5 µL PCR buffer (10 x), 0.75 µL MgCl_2_ (50mM), 1 µL of dNTPs (10mM), 1.0 µL each forward and reverse primers (10 µM), 0.5 µL *Taq* polymerase (5 U µl^−1^, Invitrogen, United States) and 5 µL template DNA (50 ng µL^−1^) in a total volume of 50 µL. The PCR amplifications were performed using a PCR thermocycler (Bio-Rad, USA). The amplified products were examined via agarose gel electrophoresis (1%) supplemented with Midori green (GC biotech, Netherlands) dye.

**Table 2.**
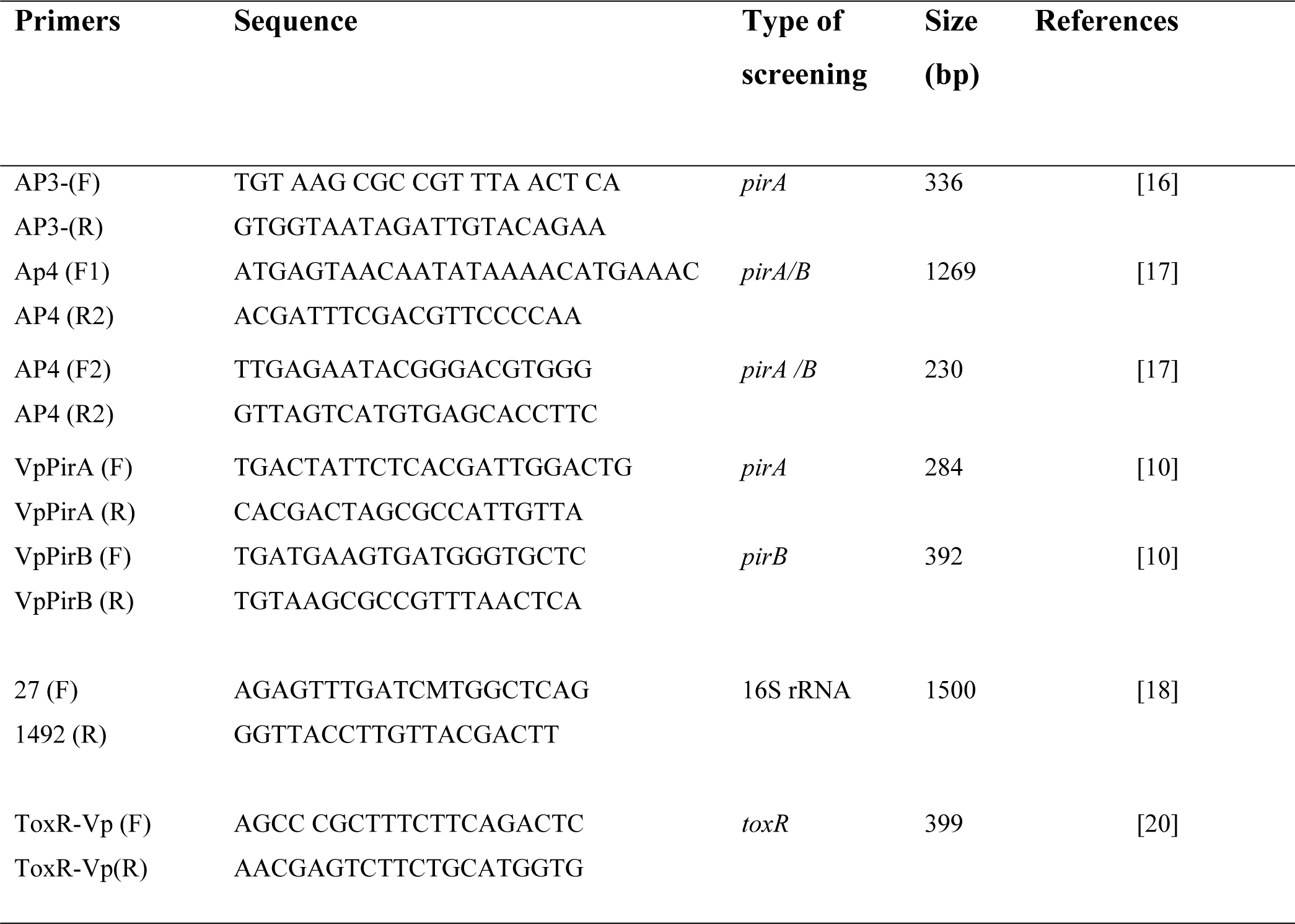
Primers used in this study

### Identification of strain NBP

The isolate was identified based on sequencing of the 16S rRNA. The primers applied for PCR amplification are listed in Table 2. The PCR mixtures were composed of the same components as mentioned above. The conditions for 16S rRNA gene amplification were (i) 4 min at 94 °C, (ii) 30 cycles of 1 min at 94 °C, (iii) 1 min 30 s at 53 °C, and (iv) 2 min at 72 °C. The amplified products were examined via agarose gel electrophoresis (1%) supplemented with Midori green (GC biotech, Netherlands) dye and were sequenced.

Bayesian analysis was performed separately for the 16S rRNA datasets using MrBayes (v3.2) [20]. A mixed model was employed for nucleotide substitutions to sample across the GTR model space. Heterogeneity rates across sites were modelled using gamma distribution. Three independent analyses with 4 Markov chains were run for 10 million generations for each data set, hence saving tree for every 1000 generations. The first 25% of the trees were removed as burn-in. The maximum clade credibility (MCC) tree of the sampled trees in Bayesian MCMC analysis and posterior probabilities (PP) of the clade have been summarised in TreeAnnotator [21]. Posterior probabilities exceeding 0.6 are represented. The evolutionary distances were computed by using the mixed model method and 0.2 nucleotide substitutions per site.

The electropherogram generated by automated DNA sequencer was read by BioEdit Sequence Alignment Editor v7.2.6.1 (http://www.mbio.ncsu.edu/BioEdit/bioedit.html) [22], wherein the sequences were carefully examined to detect missed calls and base spacing. The consensus sequences of the strains were compared with the corresponding sequences in GenBank database using Basic Local Alignment Search Tool program (BLAST; http://www.ncbi.nlm.nih.gov/BLAST/Blast.cgi). The dataset for 16S rDNA was aligned in MAFFT (v7.365) [23], and the aligned sequences were manually corrected via BioEdit. Characters that were aligned ambiguously had been excluded from the analysis. *Roseivirgo echinicomitans* strain KMM6058 (GenBank accession no. **NR043168**) was used as outgroup for 16S rDNA phylogeny.

### Co-culture of *V. parahaemolyticus* BpShHep31 and strain NBP

*Vibrio parahaemolyticus* strain BpShHep31 (at 2 × 10^3^ CFU mL^−1^) and *Algoriphagus* sp. strain NBP (at 2 × 10^5^ CFU mL^−1^) were co-cultured (cell densities 1:100 ratio respectively) at three temperatures (20 °C, 30 °C, and 40 °C). Samples were taken after 24 h, 48 h, and 72 h of co-culture. Samples were serially diluted and 100 µL aliquots of the diluted samples (dilutions 10^−6^ to 10^−9^) were plated on MA. The plates were incubated overnight at room temperature. Three colonies were picked based on their morphology and colony colour on the agar plate for screening of the presence of *pirA* and *pirB*. *Algoriphagus* sp. strain NBP colonies were re-identified after co-culture using 16S rDNA sequencing for further verification.

### Determination of conjugation efficiency of pirAB in *Algoriphagus* sp. strain NBP colonies upon co-culture

The *Algoriphagus* sp. NBP colonies that were picked after plating of the coculture with *V. parahaemolyticus* strain BpShHep31 were verified to be free from contamination by *V. parahaemolyticus* by performing PCR with specific primers for the *V. parahaemolyticus toxR* gene (which is absent in *Algoriphagus*) **(Table 2).** Sterile 10 µL pipette tip was used to pick the colonies to avoid any contamination. The DNA of the colonies were extracted by adding 10 µL of sterile distilled water followed by heating at 94°C for 5 minutes. Then, the samples were cooled at 4°C and centrifuged at 10,000 rpm for 5 minutes. The supernatant of the samples was transferred to another sterile 0.2 mL tube. The *Algoriphagus* sp. strain NBP colonies were screened for the presence of the *pirAB* genes using the VpPirA and VpPirB primers (Table 2). We then calculated the conjugation efficiency (%) as follows:

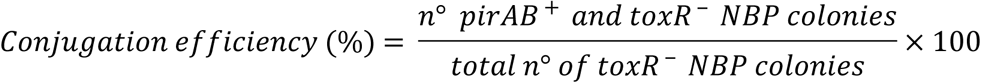

After the coculture, few colonies of *Algoriphagus* sp. strain NBP were picked from the plate and inoculated into MB and incubated overnight. Then, the culture was serially diluted and 100 µL of the aliquots of the diluted samples were plated on MA and thiosulfate citrate bile salts sucrose agar (TCBS).

### Shrimp challenge test

Healthy *Penaeus vannamei* shrimps (approximately 0.5 gram) were acclimatised for a week. The shrimps (30 shrimps in each challenge test) were transferred individually to 5 L aquariums filled with 3 L of autoclaved seawater at 25 ‰, 27 ± 1°C, 7.3 ± 0.6 mg/L dissolved oxygen, and 7.5 ± 0.6 pH. Hepatopancreases of 10 arbitrarily selected shrimps were dissected and screened to determine the presence of the *pirA* and *pirB* genes using the AP3, AP4, VpPirA and VpPirB primers (Table 2) prior to the challenge test. The isolate NBP was cultured overnight in MB for the challenge test. Hundred microlitres of the culture was spread on thiosulfate citrate bile salts sucrose agar (TCBS) to check for contamination of *V. parahaemolyticus* and incubated overnight at 28°C. Only the batch culture with negative growth on TCBS agar (used selectively to identify the presence of vibrios) and positive for *pirAB* screening used in this immersion challenge test. The immersion challenge test was carried out in triplicates through inoculation of 10^6^ CFU mL^−1^ of bacteria as described earlier [24]. Control shrimps were not exposed to any added strain. The survival rate of the shrimps was recorded daily.

The study was conducted following the Code of Practice for Care and Use of Animals for Scientific Purposes, Universiti Putra Malaysia (UPM). The study was reviewed and approved by the Institutional Animal Care and Use Committee (IACUC), Faculty of Veterinary Medicine, Universiti Putra Malaysia (UPM).

### Histopathological analysis and screening for the presence of *pirA* and *pirB* genes in challenged shrimp

Shrimps were collected on day 14^th^ (47% of survival) and immediately fixed in 10% (v/v) phosphate buffered formalin for 24 hours. After that, the shrimps were preserved in 70% ethanol until further processing. The preserved samples were sent to the Veterinary Histopathology Lab (VHL) of the Universiti Putra Malaysia (UPM) for tissue embedding and hematoxylin and eosin (H&E) staining. The hepatopancreases of five challenged and unchallenged shrimps were pulled together and screened for the presence of *pirA* and *pirB* genes using specific primers as mentioned earlier.

### Statistical analysis

The mortality data of shrimp challenged with the test bacteria, *Algoriphagus* sp. strain NBP colony count and conjugation efficiency (n°) were subjected to one-way ANOVA followed by Tukey’s post-hoc test, after prior confirmation of normality and homoscedasticity. All data are presented as mean ± standard deviation and statistical analyses were performed using SPSS version 22.

## Acknowledgements

This study was funded by Universiti Putra Malaysia High Impact Grant (vot no: 9598400). It was also supported by ‘Higher Institution Centre of Excellence’ (HICoE) grant awarded to the Institute of Bioscience (IBS), Universiti Putra Malaysia (UPM) and Japan Science and Technology Agency (JST/Japan International Cooperation Agency (JICA) through their Science and Technology Research Partnership for Sustainable Development (SATREPS-COSMOS) program with matching funds from Ministry of Education (MOE), Malaysia.

